# Fecal Microbiota Transplantation Drives Colonic Expression of Immune Activation Genes in a Mouse Model of Antibiotic Use

**DOI:** 10.1101/2021.07.23.453497

**Authors:** G. Brett Moreau, Hale Ozbek, Pankaj Kumar, Alyse Frisbee, Jhansi Leslie, William A. Petri

## Abstract

*Clostridioides difficile* infection (CDI) is the leading hospital acquired infection in North America. While the standard treatment for CDI remains antibiotics, fecal microbiota transplantation (FMT) has gained attention as an effective therapy to prevent relapse. Previous work has focused on colonization resistance mounted against *C. difficile* by FMT-delivered commensals, but the effects of FMT on the gut mucosal immune response are poorly understood. Better understanding of the molecular mechanisms driven by FMT would allow for more targeted therapy against CDI. To address this important gap in knowledge, microbial community structure and host gene expression were assessed after FMT in a mouse model of antibiotic use. Administration of FMT led to a significant increase in microbial diversity and partial restoration of community structure within 48 hours of treatment. RNA sequencing of cecal tissue identified large changes in gene expression between FMT recipient and vehicle control groups. Strikingly, genes upregulated after FMT treatment were enriched in immune activation pathways, many of which were associated with pro-inflammatory immune responses. FMT also upregulated several genes associated with type 2 immunity while repressing several associated with type 3 immunity, trends that are associated with improved response to CDI. These results highlight the interplay between the intestinal microbiota and host transcriptome and identify pathways of interest for exploring the role of FMT on treatment of recurring CDI.

## Introduction

*Clostridioides difficile* is a Gram-positive spore-forming obligate anaerobe that causes mild to severe diarrheal disease. In the CDC’s 2019 *Antibiotic Resistance Threats Report*, *C. difficile* was ranked as an urgent threat to human health (1). The increased prominence of *C. difficile* infection (CDI) is due in part to the emergence in 2005 of hypervirulent strains (2), which cause more severe disease and longer hospital stays (3–5). In a single year, *C. difficile* was associated with nearly half a million infections, 83,000 recurrences and 29,000 deaths in the United States alone (6). While *C. difficile* is one of the most common hospital-acquired infections (7), community-acquired infections are also becoming more prevalent (8). Due to the significant health impact and cost of CDI, development of highly effective treatments is critical. Risk of CDI is associated with disruption of the intestinal microbiota through antibiotic exposure (9, 10), which allows *C. difficile* to bloom and produce toxins, resulting in initiation of diarrhea, damage to the epithelium, and disruption of the intestinal barrier (7). Thus, maintaining or restoring microbiota homeostasis is critical for minimizing the risk of CDI.

While the standard treatment for CDI remains antibiotics (11), one in five patients experience recurrence (12), and antibiotic treatment further disrupts the intestinal microbiota, increasing susceptibility for relapse. Because of this, fecal microbiota transplantation (FMT), the administration of the whole microbial community from the stool of a healthy donor into the patient’s gastrointestinal tract, has been explored as a therapeutic for CDI. While FMT has been implicated as therapeutic in a wide range of disorders (13), it has proved most successful as treatment for CDI, with an efficacy of over 90% (14). Despite the efficacy of FMT at treating CDI, it is still considered an experimental therapy due to concerns of FMT donor standardization, the evolving regulation of FMT, long-term side effects, and risk of pathogen transmission, particularly for immunocompromised patients (15). FMT has been associated with transmission of antibiotic resistant pathogens to recipients, including one fatality (16). Given the prevalence of recurrent CDI and the potential risks posed by FMT, more defined therapeutics for CDI are necessary. Better understanding of the protective mechanisms behind FMT will allow for more targeted therapeutics such as a defined consortium of probiotic organisms or a defined cocktail of immunotherapeutic drugs as a replacement for FMT.

Because *C. difficile* disease is initiated by antibiotic disruption of the intestinal microbiota (9, 10), much of the focus of FMT has been on its ability to provide colonization resistance against *C. difficile* (17). Recurrent CDI is characterized by decreased microbial diversity that is associated with depletion of the *Bacteroidota* and *Firmicutes* phyla and spikes in *Proteobacteria* (18), and these changes in diversity can be restored by FMT (19). FMT may also work by influencing the immune response in the gut to facilitate *C. difficile* clearance and prevent excessive host damage. It has recently been more greatly appreciated that the type of host immune response can impact CDI severity (20). While *C. difficile* toxins cause much of the virulence and epithelial barrier damage seen in CDI, an especially robust immune response and resulting inflammation has been implicated in severe disease and worse clinical outcome independently of bacterial burden (21–23). Specific types of immune responses have also been implicated in differential CDI outcomes: tissue-regulated type 2 immunity has been associated with tissue repair and protection from severe disease (24, 25), while a type 3 immune response has been implicated in increased disease severity (26). Balancing these immune types to produce a response that facilitates *C. difficile* clearance while minimizing tissue damage is critical for decreased disease severity. As components from the microbiota and microbially-derived metabolites can influence immune responses (20), this may be another mechanism through which FMT drives resistance to CDI.

More mechanistic understanding of FMT’s impact on the gut is required to develop more defined therapeutics. While the ability of FMT to increase microbial diversity and restore microbial community composition has been well studied (27–30), its effect on the host transcriptome remains poorly understood. Further limiting our understanding of FMT is that many studies have investigated FMT in the context of other intestinal disorders, such as intestinal colitis models (30–32). In addition, studies often look well after FMT administration (33), making it difficult to distinguish the effects of FMT from the effects of gradual microbiota recovery in the absence of antibiotics. This study aims to address these gaps in knowledge by investigating the immediate impact of FMT in the context of antibiotic depletion of the host microbiota. The primary findings of this study demonstrate that FMT elicited significant changes in microbial composition and gene expression within 48 hours of administration and that changes in gene expression were enriched in host immune activation gene sets. In particular, changes to immune genes suggest an upregulation of pro-inflammatory and resolving type 2 immune signatures while suppressing type 3 activation. This work highlights the crosstalk between the intestinal microbiota and immune responses in the gut, identifying a potential mechanism by which FMT could protect against CDI.

## Results

### FMT Partially Restores Microbiome Diversity after Antibiotic Treatment

To assess the effect of FMT on microbiome composition and host gene expression, mice were given antibiotics to disrupt the gut microbiota, then treated with either FMT or vehicle control for two days. Forty-eight hours after the initial FMT administration, FMT recipient, vehicle control, and FMT donor mice were sacrificed (Figure 1). Cecal contents and cecal tissue samples were collected for analysis of microbiota community structure and host gene expression, respectively. First, the V4-16S rRNA gene was sequenced using DNA from cecal contents to examine microbial community structure. Analysis identified 566 unique amplicon sequence variants (ASVs) in these samples. Community diversity in each group was assessed with the hypothesis that antibiotic treatment would decrease community diversity and that diversity would be restored by FMT.

**Figure 1:**
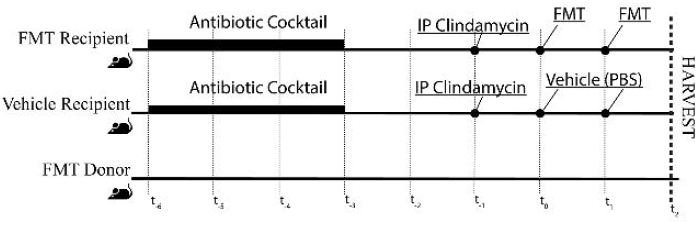
*Overview of experimental design.* C57BL6 mice were divided into three groups: FMT donors (n=12), FMT recipients (n=10), and vehicle controls (n=10). FMT recipient and vehicle control mice were given an antibiotic cocktail in drinking water for three days, then switched to regular drinking water and given a single IP injection of Clindamycin on Day −1. On Day 0 and Day 1, mice were administered either FMT (a fecal slurry from donor mice) or vehicle (PBS) by oral gavage. On Day 2, cecal contents and cecal tissue were harvested from mice for downstream analysis. FMT donor mice were untreated until harvest on Day 2.

The alpha diversity of the microbial community was first assessed by looking at both the overall ASVs observed as a measure of sample richness as well as Shannon’s diversity index, which accounts for both richness and evenness. We observed significant differences between all groups (Figure 2A) as measured by a pairwise Wilcoxon Rank test with Bonferroni correction for multiple comparisons. Treatment with the antibiotic cocktail was sufficient to disrupt diversity in the vehicle control group relative to the FMT donor group, which was not treated with antibiotics. While FMT administration partially restored diversity, it was unable to reach the level observed in the FMT donor group.

**Figure 2:**
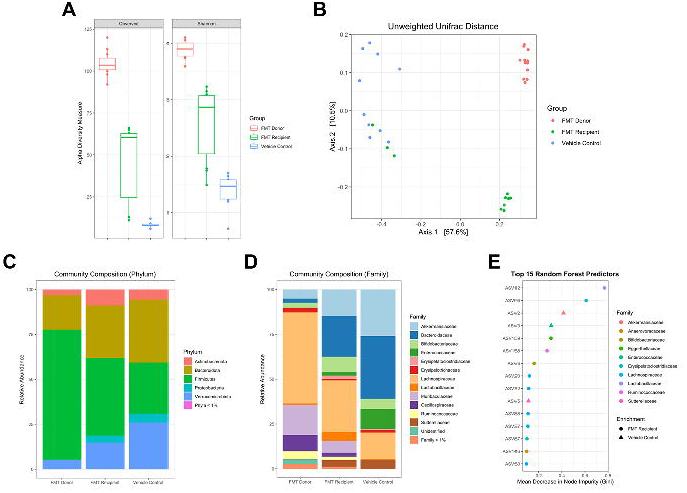
*FMT partially restores the microbiome after antibiotic disruption.* A) Alpha diversity metrics (Total number of observed ASVs and Shannon Index) for FMT donor (red), FMT recipient (green) and vehicle control (blue) groups. All groups within each metric are significantly different from each other when analyzed using a pairwise Wilcoxon Rank test with Bonferroni correction for multiple comparisons. B) Non-metric multidimensional scaling (NMDS) plot of beta diversity (unweighted Unifrac) metrics for FMT donor (red), FMT recipient (green) and vehicle control (blue) groups. The percent of variance explained by each dimension is listed on each axis. C-D) Relative abundance of taxa within each group at the C) phylum and D) family levels. Taxa that comprised less than 1% of relative abundance in all groups are aggregated. E) Ranking of the Top 15 ASVs that distinguish FMT recipient samples from vehicle controls as determined by a random forest model. ASVs that are more abundant in the FMT recipient group relative to vehicle controls are displayed as circles, while ASVs more abundant in the control group are displayed as triangles.

Beta diversity between samples was also assessed using Unweighted Unifrac distance, which takes into account phylogenetic distance between ASVs. Similar to the results from alpha diversity metrics, FMT donor, vehicle control, and FMT recipient groups all clustered separately. In addition, the FMT recipient group clustered more closely with the FMT donor group along Axis 1, which accounted for most of the variation in the model (Figure 2B). Several FMT recipient samples clustered with the vehicle control group, and these same samples also had lower alpha diversity values more similar to those seen in control samples. It is possible that the FMT failed to establish in these animals, which may account for why they are more similar to the vehicle control than other FMT recipient samples. Similar results were observed using Bray-Curtis Dissimilarity index as a different measure of beta diversity (Supplemental Figure 1), indicating that this outcome is not dependent on bacterial phylogeny. These data indicate that FMT administration partially restored community diversity to levels more similar to those seen in mice that did not receive antibiotics.

### FMT Restores Community Composition by Increasing Relative Abundance of ***Firmicutes***

Next, community composition was assessed to identify relative changes at the phylum and family levels between groups. The FMT donor group was composed primarily of members of the *Firmicutes* phylum, with the *Bacteroidota* phylum making up a minor proportion (Figure 2C) The *Firmicutes* phylum was a smaller proportion of the antibiotic-treated vehicle control samples relative to the FMT donor group, with antibiotic-treated mice having relatively higher abundance of *Verrucomicrobiota*, *Bacteroidota*, and *Proteobacteria.* FMT administration after antibiotic treatment restored a community composition more closely resembling the FMT donor group, with increased *Firmicutes* and decreased *Bacteroidota*, *Verrumicrobiota*, and *Proteobacteria* relative to the vehicle control group.

Similar differences in community composition were observed at the Family level, with the relative abundance of multiple families being intermediate in the FMT recipient group between the FMT donor and vehicle control groups (Figure 2D). The most prominent families observed in the FMT donor group were *Lachnospiraceae*, *Muribaculaceae*, and *Oscillospiraceae*, whereas these families made up a minority of the vehicle control samples in favor of relative increases in the proportion of *Akkermansiaceae*, *Bacteroidaceae*, and *Enterococcaceae*. Relative to the vehicle control, mice that received FMT had increases in the proportion of *Lachnospiraceae*, *Muribaculaceae*, and *Oscillospiracea* as well as decreases in *Akkermansiaceae*, *Bacteroidaceae*, and *Enterococcaceae*, leading them to a more similar community composition to that seen in the FMT donor group. Overall, these data indicate that administration of FMT partially restored the microbial community in antibiotic-treated mice to a similar composition to that of the FMT donor.

Finally, a random forest model was generated to identify ASVs that best discriminated between vehicle control and FMT recipient groups. The model performed well, with a 14% out-of-bag error rate. Of note, the model correctly classified all samples except for the three FMT recipient samples that exhibited lower diversity and clustered with Vehicle control samples. Of the top 15 ASVs selected by the random forest model, 87% (13/15) were more prevalent in the FMT recipient group relative to controls (Figure 2E), with most of these ASVs from the *Firmicutes* phylum. At the genus level, FMT recipient samples had increased relative abundance of *Lactobacillus ssp.* and *Erysipelatoclostridium ssp.*, while *Akkermansia muciniphilia* and *Enterococcus ssp.* had higher relative abundance in the control samples relative to FMT recipients. The top 15 ASVs were also enriched in ASVs from the *Lachnospiraceae* family, which were largely absent from vehicle control samples but present in the FMT recipient group. These data highlight the significance of restoration of the *Firmicutes* phylum after FMT.

### FMT Significantly Changes the Host Transcriptome

The immediate effect of FMT on host gene expression was next assessed. Cecal contents from a subset of mice (four mice/group) were selected for whole RNA sequencing. Of note, all four FMT recipient samples came from mice that displayed increased alpha and beta diversity metrics relative to vehicle controls. Community diversity and composition analysis in this subset of samples was representative of diversity and composition observed in the full data set (Supplemental Figure 2).

Principal component analysis (PCA) was first performed on normalized, variance-stabilized and log_2_-transformed count data to further check data quality and assess similarity between groups. FMT donor samples clustered separately from vehicle control and FMT recipient samples along PC2, which likely indicates the effect of antibiotic treatment on transcription (Figure 3A). While there was overlap between vehicle control and FMT recipient samples, FMT recipient samples appeared to cluster more closely with FMT donor samples along PC1, which accounted for most of the total variation in the PCA model. These results suggest that the most prominent factor distinguishing RNA sequencing results from different samples was antibiotic treatment. While differences in host gene expression were less pronounced between FMT recipient and vehicle control samples than differences in community diversity, there appeared to be some clustering between these samples, with the FMT recipient group appearing more similar to FMT donors.

**Figure 3:**
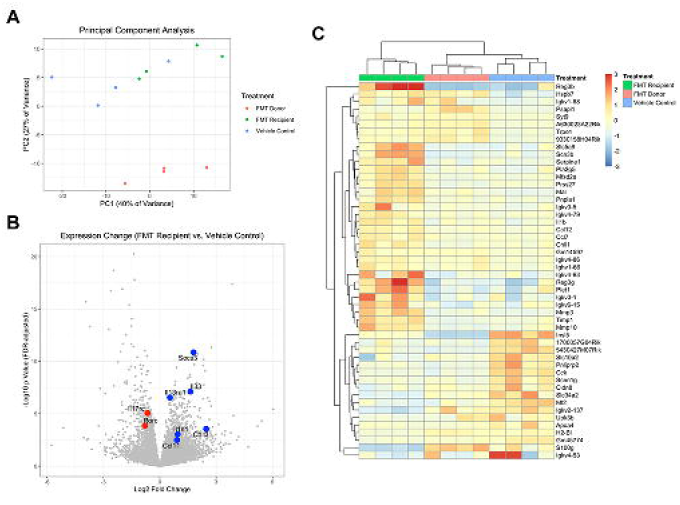
*FMT significantly changes host gene expression.* A) Principal component analysis of RNA sequencing reads from FMT Donor (red), FMT recipient (green), and vehicle Control (blue) samples. B) Transcriptional changes in FMT recipient group relative the vehicle control group. Select type 2 immunity-associated genes are colored in blue, while select type 3 immunity-associated genes are colored in red. *SOCS3* is both a promoter of type 2 responses and an inhibitor of type 17 responses. C) Heatmap of normalized and log2-transformed gene counts for genes with at least a four-fold change in expression and FDR-adjusted p-value < 0.05 between FMT recipient and vehicle control samples. Samples and genes are both hierarchically clustered.

Previous work from our lab has demonstrated that type 2 immune responses, particularly the alarmins IL-25 and IL-33, are protective against CDI, in part by activating group 2 innate lymphoid cells (ILC2) and promoting an eosinophilic response at the site of infection (24, 25). Based on these data, a directed approach was taken to evaluate expression of type 2 genes with the hypothesis that FMT would promote their expression. While *IL25* expression was not detected in these samples, expression of *IL33* and its receptor ST2 (*IL1RL1*) was upregulated in response to FMT (Figure 3B). In addition, expression of the major eosinophil attractant Eotaxin (*CCL11*) trended higher (FDR-adjusted p = 0.056) in FMT treated mice, consistent with FMT promoting an immune response that facilitates eosinophil recruitment. Other markers of type 2 immune responses including YM1 (*CHIL3*) (34, 35) and the IL-13 receptor (*IL13RA1*) were also upregulated in response to FMT. The ability of FMT to repress IL-17A and related type 3 immune responses was also of interest, as induction of a type 3 immune response can promote more severe disease during CDI (26). While *IL17* was not detected, expression of the IL-17 co-receptor (*IL17RC*) and the transcription factor RORγt (*RORC*) were inhibited in response to FMT. *SOCS3*, a signaling regulator that promotes type 1 or 2 immune responses while repressing type 3 differentiation (36), was also upregulated in response to FMT. Overall, these findings support the hypothesis that FMT administration may promote shifts towards a type 2-mediated immune response and away from a type 3 response.

### RNA Sequencing Implicates Inflammatory Biomarkers and Altered Lipid Metabolism in the Immediate Response to FMT

Next, an unbiased approach was taken to examine transcriptional differences between vehicle control and FMT recipient samples. A total of 864 genes were differentially expressed between vehicle control and FMT recipient samples after false discovery (FDR) correction (Supplemental Table 1). Of these genes, 292 had at least a 2-fold change (log_2_ fold change > 1.0) expression between groups and 50 had at least a 4-fold change (log2 fold change > 2.0). Of these 50 genes, 33 were upregulated in the FMT group relative to the vehicle control and 17 were downregulated in the FMT condition. Unsupervised hierarchical clustering was performed on these 50 genes to identify patterns within this gene set (Figure 3C). Samples clustered according to group, indicating that all three groups had unique expression patterns.

Many of the upregulated genes in the FMT recipient group encode proteins associated with the host immune response. These include chemokines *CCL7*, *CCL12*, and *IL1B* as well as immune signaling pathway proteins, such as the Toll adapter protein *MAL* (37) and *PLET1*, which facilitates dendritic cell migration into the intestine (38). Biomarkers of intestinal inflammation such as the Chitinase 1 (*CHIT1*) gene and inflammasome-dependent antimicrobial peptides *REG3B* and *REG3G* (39) were also upregulated in the FMT recipient group compared to vehicle controls. Immunoglobulin kappa variable genes were identified as both upregulated and downregulated in response to FMT, but were more abundant in the FMT group compared to controls. Extracellular matrix remodeling was also enriched in response to FMT, with upregulated expression of both matrix metalloproteinases (*MMP3*, *MMP10*, *PRSS27*) and proteinase inhibitors (*TIMP1*, *SERPINE1*), pathways that are expressed in response to immune signaling (40).

Downregulated genes made up a minority (17/50, 34%) of differentially expressed genes and were more heterogeneous in their functions. Genes encoding immune-associated proteins were also downregulated in response to FMT, including *S100G*, Metallothionein 2 (*MT2*), and the tight junction protein Claudin 8 (*CLDN8*). Several genes encoding bile acid and ion transporters were also downregulated after FMT, including *SLC34A2*, *SLC10A2*, and *SCNN1G*. Finally, several genes were involved in host metabolism, such as the Apoliprotein A-IV *APOA4*, the pancreatic lipase *PLRP2*, and the peptide hormone *CCK*. Overall, genes with the largest changes in gene expression were consistent with increased immune activation in response to FMT.

### GSEA-Assisted Pathway Analysis Identifies Immune Activation and Decreased Lipid Processing in Response to FMT

To better understand the biological processes broadly associated with FMT, Gene Set Enrichment Analysis (GSEA) (41, 42) was performed utilizing the entire gene expression data set. Overall, 23 gene sets had an FDR-adjusted p < 0.05 value for significance (Figure 4), with 74% (17/23) being upregulated in FMT recipient samples relative to vehicle controls. Upregulated gene sets were dominated by immune pathways, including “HALLMARK_INFLAMMATORY_RESPONSE”, “HALLMARK_IL6_JAK_STAT3_SIGNALING”, and “HALLMARK_INTERFERON_GAMMA_RESPONSE”, suggesting an immediate pro-inflammatory response to FMT administration. Many genes were associated with multiple immune gene sets, with core genes such as *CCL2*, *SOCS3*, and *TIMP1.* There was also significant overlap with genes previously identified in Figure 3C, such as *SERPINE1, CCL7*, and *IL1B*. In addition to these pathways, the gene set “HALLMARK_EPITHELIAL_MESENCHYMAL_TRANSITION” was upregulated after FMT, which may suggest genes involved in resolution of inflammation are being upregulated.

**Figure 4:**
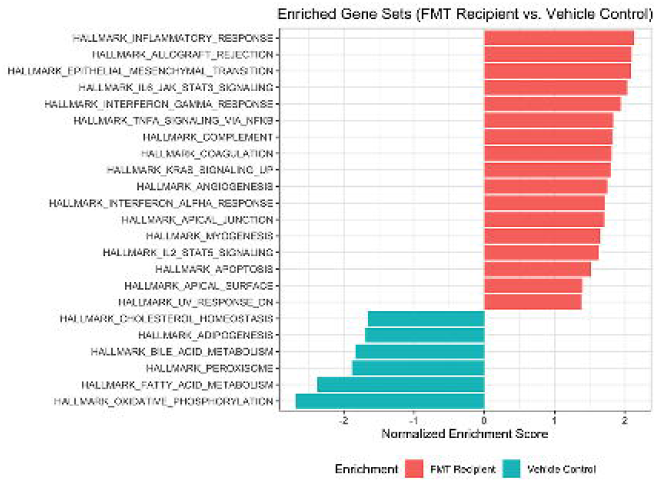
*FMT increases expression of host immune pathways.* Ranking of significantly enriched (FDR adjusted p-value < 0.05) gene sets in either FMT recipient (red) or vehicle control (blue) samples. Gene sets are ranked according to Normalized Enrichment Score, which normalizes gene enrichment based on the size of each gene set.

Downregulated gene sets primarily consisted of metabolism pathways, including “HALLMARK_OXIDATIVE_PHOSPHORYLATION”, “HALLMARK_FATTY_ACID_METABOLISM”, and “HALLMARK_BILE_ACID_METABOLISM”. The “HALLMARK_OXIDATIVE_PHOSPHORYLATION” gene set was distinct from other downregulated gene sets, with notable genes enriched in this data set encoding proteins such as Monoamine Oxidase B (*MOAB*), Pyruvate Dehydrogenase Kinase 4 (*PDK4*), ATPase Na+/K+ transporting subunit beta 1 (*ATP1B1*), and Cytochrome C Oxidase Copper Chaperone 11 (*COX11*). There was significant overlap between genes in the remaining gene sets. Core genes associated with these pathways encode proteins involved in fatty acid synthesis, such as Acyl-CoA Synthetase (*ACSL1*), Malonyl-CoA Decarboxylase (*MLYCD*), and Aldehyde Dehydrogenase 1 (*ALDH1A1*). Overall, these results suggest that FMT is associated with broad changes in immune activation as well as downregulation of metabolic pathways.

## Discussion

While the effect of FMT on the intestinal microbiota has been well established, its impact on host gene expression is more poorly understood. To address this gap in knowledge, mice were treated with antibiotics before administration of either FMT or vehicle control. Overall, the primary finding from this work demonstrates that FMT results in significant changes to the host transcriptome relative to antibiotic treatment alone, and that these genes are enriched in immune activation pathways. Immune genes, including both pro-inflammatory pathways and several genes associated type 2 immunity were upregulated in response to FMT treatment. In contrast, several pathways were downregulated after FMT, including oxidative phosphorylation, fatty acid metabolism, and bile acid metabolism. In addition, this study confirms previous findings that FMT significantly increases microbial diversity and partially restores community composition to that seen before antibiotic treatment.

Microbiome diversity and community structure were both partially restored in antibiotic treated mice within 48 hours of initial FMT administration. This finding is consistent with previous studies that observed effective restoration of microbiome diversity and structure after FMT treatment in both animal models (28, 30) and clinical studies (27, 29, 43, 44). Restoration of *Firmicutes* was a specific hallmark of FMT treatment, as 13 of the 15 ASVs that best distinguish FMT-treated mice from vehicle controls were from this phylum. Notable *Firmicutes* restored by FMT include *Lactabacillus ssp.*, which was the ASV that most strongly predicted FMT treatment, as well as multiple taxa from the *Lachnospiraceae* family. These changes suggest a microbial environment post-FMT that is more resistant to CDI, as FMT success at preventing recurrent *C. difficile* infection is associated with restoration of the *Firmicutes* and *Bacteroidota* phyla (45). In particular, several *Lactobacillus* species have been implicated in protection against recurrent CDI (46), potentially through production of organic acids that inhibit *C. difficile* growth or inhibition of *C. difficile* toxin (47). Similarly, *Lachnospiraceae* species provide colonization resistance against *C. difficile* in a mouse model, leading to decreased intestinal histopathology and improved survival (48). This may be due in part to production of short-chain fatty acids from carbohydrates, which inhibit *C. difficile* growth, mediate epithelial proliferation, promote barrier integrity, and exert anti-inflammatory effects in the gut (49, 50). In contrast, the three taxa enriched in the control group relative to FMT-treated samples were *Akkermansia mucinophilia* from the *Verrucomicrobiota* phylum, *Enterococcus ssp.* from the *Firmicutes* phylum, and *Sutterellaceae ssp.* from the *Proteobacteria* phylum. Increased relative abundance of the *Proteobacteria* phylum and *Enterococcus* family have been observed immediately after antibiotic treatment (51), and these taxa are reduced after FMT treatment (52). Disruption of the microbiota resulting in increased relative abundance of *Proteobacteria* and *Enterococcus* also triggered *C. difficile* overgrowth and increased spore shedding (53), suggesting that increased relative abundance of these taxa may promote CDI. Taxa of the phylum *Verrucomicrobiota* (including *Akkermansia*) are enriched after broad-spectrum antibiotic treatment (54), indicating that higher relative abundance of this taxa in the vehicle control group may be due to antibiotic treatment. Overall, these results are largely consistent with other findings in the field and suggest that FMT administration restores a microbial community more similar to before antibiotics and more resistant to *C. difficile*.

A key finding of this work is the significant impact of FMT administration on host gene expression. A total of 864 genes were differentially expressed between FMT treated and vehicle control samples, with a significant proportion of these genes involved in immune activation pathways. In particular, GSEA analysis identified an enrichment in immune pathways that are primarily considered to be pro-inflammatory, such as IL-6, IFNγ, and TNFα. There is some evidence that disruption of the intestinal microbiota is associated with disruption of type 1 immune responses and that these responses can be restored through administration of FMT or microbial associated molecular patterns (MAMPs) (55, 56). Therefore, the introduction of MAMPs through FMT may drive the enrichment in immune genes observed in this study. Activation of these immune pathways may facilitate some of the protective effects of FMT independently of the microbiota, as previous work identified filtered fecal supernatants as sufficient to protect against recurrent CDI (57). Previous studies have found that FMT diminishes production of pro-inflammatory cytokines such as IL-6 and TNFα (33) while promoting anti-inflammatory mediators and regulatory T cells (30, 58). However, many of these studies were in the context of intestinal inflammatory disorders and looked later in the course of FMT than the current study, so it is unclear how comparable they are to the acute phase of FMT in a naive setting. Strong inflammatory responses in the initial response to infection are critical for *C. difficile* clearance and CDI recovery (59–61), and it is possible that the upregulation of inflammatory genes immediately after FMT promotes a stronger response upon subsequent *C. difficile* infection. Additional studies in this context are necessary to elucidate the impact of these immediate pro-inflammatory responses on CDI.

Signatures of type 2 immune responses and wound repair pathways were also observed after FMT administration, indicating that the microbiota plays a key role in maintaining a tissue regulatory and reparative response. Directed analysis of type 2 and type 3 associated genes in the list of differentially expressed genes identified several type 2 genes (*IL33*, *IL1RL1*, *CHIL3*, *CCL11*) that were upregulated and several type 3 genes (*RORC*, *IL17RC*) that were downregulated in response to FMT. *SOCS3*, a critical signaling molecule that promotes type 2 and inhibits type 3 differentiation, was also identified in multiple gene sets enriched after FMT. These results are consistent with previous results finding decreased expression of type 2 cytokines in germ-free mice and restoration of their expression after FMT treatment (62, 63). Specific taxa identified in this study may be critical to these shifts: for example, *REG3B* and *REG3G*, which were upregulated in response to FMT, were restored after *Lactobacillus* supplementation in a DSS colitis model (64). Type 2 immune responses are associated with wound repair and eosinophil recruitment in response to infection or inflammation (65), indicating that upregulation of these genes may be the next step in restoration of intestinal homeostasis. Indeed, genes involved in the epithelial to mesenchymal transition, a key process in wound repair after inflammatory signaling (66), were also identified after FMT by GSEA. Several genes in this set were also identified in the top 50 differentially regulated genes, including the metalloprotease *MMP3* and protease inhibitors *TIMP1* and *SERPINE1*, which are important for extracellular matrix remodeling (67, 68). In the context of CDI, type 2 responses and subsequent eosinophilic responses are critical to reduce disease severity (24, 25). Type 3 responses are slightly more mixed: early pro-inflammatory responses are required for *C. difficile* clearance, but prolonged inflammation and neutrophil recruitment is associated with worse outcomes (26, 69, 70). Together, these gene expression data are consistent with the paradigm that a type 2 immune response is generally protective against CDI while a type 3 immune response exacerbates disease, which may partially explain one of the mechanisms by which FMT protects against CDI.

Several pathways were downregulated in FMT-treated samples compared to vehicle controls, including oxidative phosphorylation, fatty acid metabolism, and bile acid metabolism. Downregulation of oxidative phosphorylation genes is consistent with observations that immune cells are activated in this condition, as immune activation is accompanied by a shift in immune cell metabolism from oxidative phosphorylation to glycolysis (71). While immune activation is likely a key driver of the downregulation of oxidative phosphorylation genes, this may also be an effect of antibiotic treatment. Oxidative phosphorylation, fatty acid metabolism, and bile acid metabolism genes were all downregulated in antibiotic treated mice (44), highlighting the influence of the intestinal microbiota on these pathways. The microbiota is known to significantly influence lipid absorption and metabolism (72), indicating that disruption of the microbiota can impact lipid profiles relative to the healthy gut. *SLC10A2*, which encodes a major primary bile acid importer, was also downregulated after FMT treatment. Expression of *SLC10A2* is downregulated by the intestinal microbiota (73), suggesting that restoration of the intestinal microbiota may prevent bile acid reabsorption and increase the pool of microbially-modified secondary bile acids, which are protective against CDI (74). Further research is necessary to determine how these changes in gene expression affect bile acid and lipid profiles in the gut and their downstream effects on disease susceptibility and severity.

Together, these data show that FMT quickly restores microbial diversity and community composition. FMT also significantly alters host gene expression, particularly immune genes, demonstrating the dynamic and highly coupled relationship between the host and the microbiota to maintain homeostasis. Many of these pathways were associated with pro-inflammatory immune responses, which may indicate an immediate response to the microbial community or microbial components in the FMT. In addition, FMT upregulated several genes associated with type 2 immune differentiation and repressed several type 3 biomarkers, suggesting a shift towards repair and resolution of inflammation. In the context of CDI, these shifts suggest an environment that is more resistant to subsequent *C. difficile* infection. Future studies are required to identify how long-lasting these transcriptional changes are and how they change the response to *C. difficile* during infection. However, these results highlight the crosstalk between the intestinal microbiota and host gene expression while identifying modulation of the immune response as an important potential mechanism through which FMT protects against intestinal dysbiosis and subsequent infection.

## Acknowledgements

This work was funded by the US National Institutes of Health (1R01 AI152477 and R01 AI124214 to WAP, 1F31AI136421-01A1 and 5T32AI007496-24 to ALF, and F32DK124048 to JL). The funders played no role in experimental design or data interpretation.

## Methods

### Murine FMT Model

All work with animals complied with all relevant ethical regulations for animal testing and research procedures approved by Institutional Animal Care and Use Committee at the University of Virginia (IACUC). Murine experiments were carried out using eight-week-old male C57BL6/6J mice (Jackson Laboratory). Mice were split into three groups: FMT donor (n=12), FMT recipient (n=10), and vehicle control (n=10). Mice in vehicle control and FMT recipient groups were given an antibiotic cocktail in drinking water consisting of 45mg/L Vancomycin (Mylan), 35 mg/L Colistin (Sigma), 35 mg/L Gentamicin (Sigma), and 215 mg/L Metronidazole (Hospira) for three days. Mice were then switched to normal drinking water and given a single IP injection (0.016 mg/g) of Clindamycin (Hospira) on Day -1.

On Day 0 and Day 1, FMT recipient mice were given an oral gavage of 100ul FMT, while vehicle control mice were gavaged with anaerobic PBS as a sham control. FMT was prepared from fecal pellets from age- and sex- matched donor mice that were collected and immediately homogenized by vortexing in 1.5ml sterile anaerobic PBS. Mice were sacrificed 48 hours after the initial FMT treatment on Day 2. Cecal contents were collected for 16S rRNA sequencing and harvested cecal tissue were immediately placed in RNA*later* (Sigma) and stored at −80°C.

### 16S rRNA Gene Sequencing

DNA extraction and 16S rRNA sequencing from murine cecal content was performed as previously described (25, 28). Briefly, DNA was extracted using the MagAttract PowerMicrobiome kit (Qiagen) with the EpMotion liquid handling system (Eppendorf). The V4 region of the 16S rRNA gene was amplified from each sample using a dual indexing sequencing strategy (75). Samples were sequenced on the MiSeq platform (Illumina) using the MiSeq Reagent Kit V2 500 cycles (Illumina #MS102-2003) according to the manufacturer’s protocol with modifications found in the Schloss Wet Lab SOP (https://github.com/SchlossLab/MiSeq_WetLab_SOP).

### 16S Sequence Analysis

All analyses were performed in R (version 4.0.2) (76). Raw sequencing reads were inspected for quality, filtered, paired reads merged, and chimeric sequences removed using DADA2 (version 1.18.0) (77). Length distribution of non-chimeric sequences was plotted to ensure lengths matched the expected V4 amplicon size. Taxonomy was assigned to amplicon sequence variants (ASVs) by aligning with the Silva reference database (version 138.1) (78). Alpha and beta diversity were calculated using the phyloseq package (version 1.34.0) (79). The packages tidyverse (version 1.3.0) (80) and ggpubr (version 0.4.0) were used for data organization and analysis.

Sample preparation and sequencing quality was assessed using blank control, water control, and mock community control samples. Blank and water control samples had significantly fewer reads than other samples, suggesting that contamination was not a major problem during the sequencing run. A mock community control sample was also processed to investigate sequencing accuracy and identified all eight community members included in the mock community sample at similar relative abundance (Supplemental Table 2). Three additional ASVs were identified, including a *Bacteroidota* ASV that comprised 0.17% of sequence abundance, a LD29 ASV that comprised 0.05% of sequence abundance, and an ASV of unknown genus that comprised 2.1% of sequence abundance. This unknown ASV is of the *Enterobacteriaceae* Family, which is the same family as *Salmonella ssp.*, a predicted member of the community. We hypothesize that this ASV of unknown genus may represent *Salmonella* that was artificially split during the DADA2 pipeline (81).

Random forest analysis was performed in R using the randomForest package (version 4.6.14) (82). Samples were first divided into training (70% of samples, divided equally between FMT recipient and vehicle control samples) and test (30% of samples) sets. The training set was used for tuning the “mtry” parameter of the model, while the test set was used to validate model performance. Feature importance was determined using the Gini index, which measures the total decrease in node impurity averaged across all trees.

### RNA Isolation from Mouse Cecal Tissue

RNA*later*-stabilized cecal tissue was removed using sterile needles and placed in 1ml TRIzol per tissue. Tissues were disrupted and homogenized using TissueLyser II according to the manufacturer’s protocol and then tubes were centrifuged to separate the aqueous and organic phases. The aqueous phase was precipitated in ethanol, and RNA was redissolved in 400ul RLT buffer and further purified using the RNeasy kit (Qiagen) according the manufacturer’s protocol. The size, distribution, and quantity of total RNA in each sample was determined using an Agilent 2100 bioanalyzer. All samples had an RNA integrity number (RIN) above 8.

### RNA Sequencing

RNA Sequencing was performed by Novogene. Briefly, mRNA was isolated using oligo(dT) beads and randomly fragmented, followed by cDNA synthesis and second-strand synthesis. After end repair, barcode ligation, and sequencing adaptor ligation, the double-stranded cDNA library was size selected and PCR amplified. Sequencing was carried out on an Illumina HiSeq platform PE150 with paired-end 150 base pair reads.

### Gene Expression Profiling

RNA sequencing libraries were checked for their quality using the FastQC program (https://www.bioinformatics.babraham.ac.uk/projects/fastqc/) and aggregated using MultiQC (83). Clean reads were mapped to the most updated version of the transcriptome with the STAR aligner, and counted using HTseq (84). Count tables were imported into R for differential gene expression analysis. The DESeq2 package (85) was used to exclude genes with low counts, normalize data, estimate dispersion, and fit counts using a negative binomial model. A heatmap of differentially expressed genes between FMT recipient and vehicle control groups was generated from normalized count data in R using the pheatmap package (version 1.0.12). Differentially expressed genes were also ranked (upregulated to downregulated) based on log_2_ fold change and FDR corrected p-values for pathway analysis using Gene Set Enrichment Analysis (GSEA) software (41, 42). Hallmark gene sets from MSigDB were chosen for this analysis.

### Data Sharing

Raw sequencing reads have been deposited in the Sequence Read Archive (SRA) database under Accession number PRJNA521980. Data and code for data processing are hosted at https://github.com/gbmoreau/FMT_Mouse_16S_RNAseq.

## Notes

### Competing Interest Statement

The authors have declared no competing interest.

### Summary of Updates

Updated to add figures at higher size/resolution.

