## Supplemental Figures (1 and 2) for "Fecal Microbiota Transplantation Drives Colonic Expression of Immune Activation Genes in a Mouse Model of Antibiotic Use"

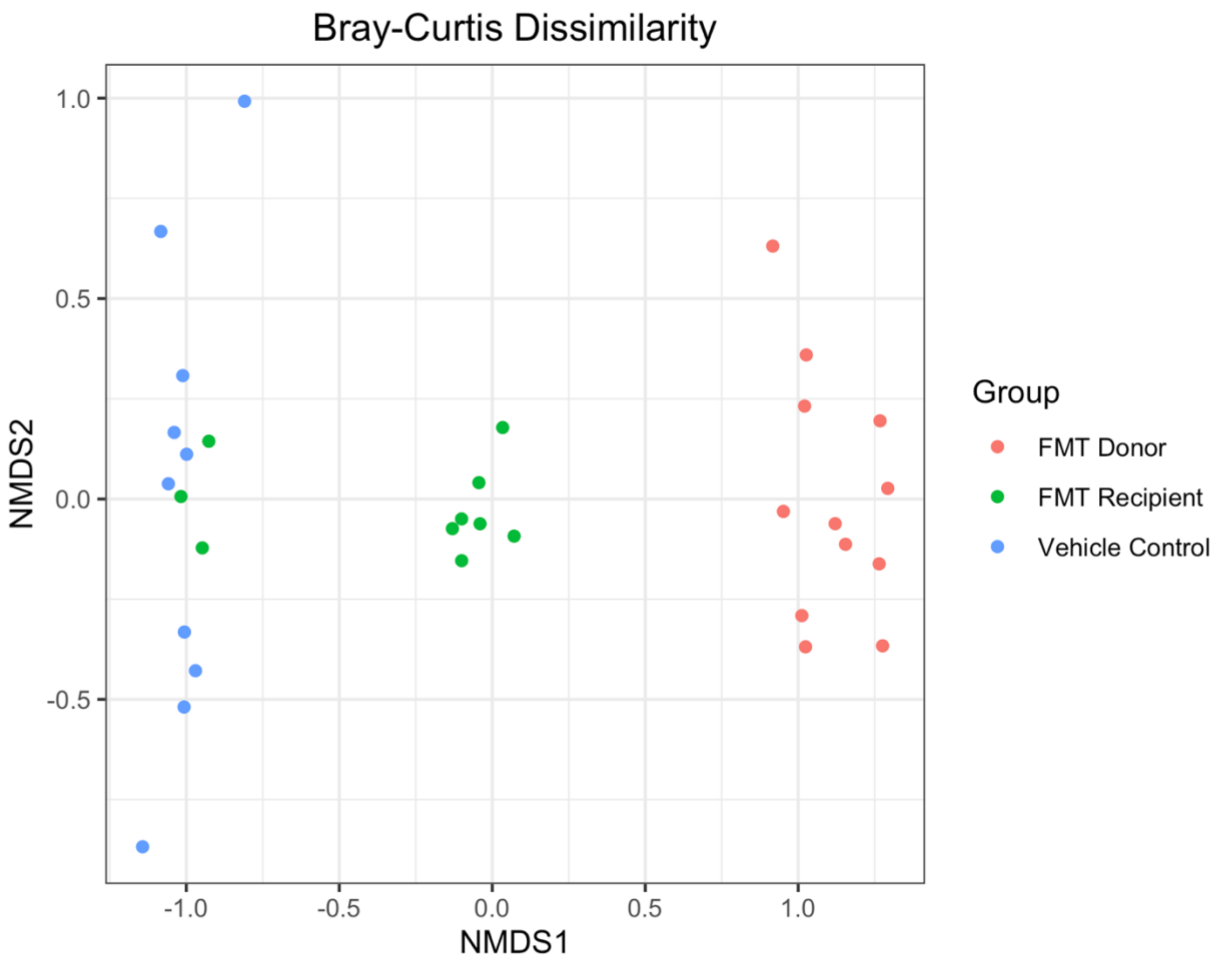


**Supplemental Figure 1:** *FMT recipient samples have intermediate beta diversity.* NMDS plot of Bray-Curtis Dissimilarity index for FMT donor (red), FMT recipient (green), and vehicle control (blue) samples.

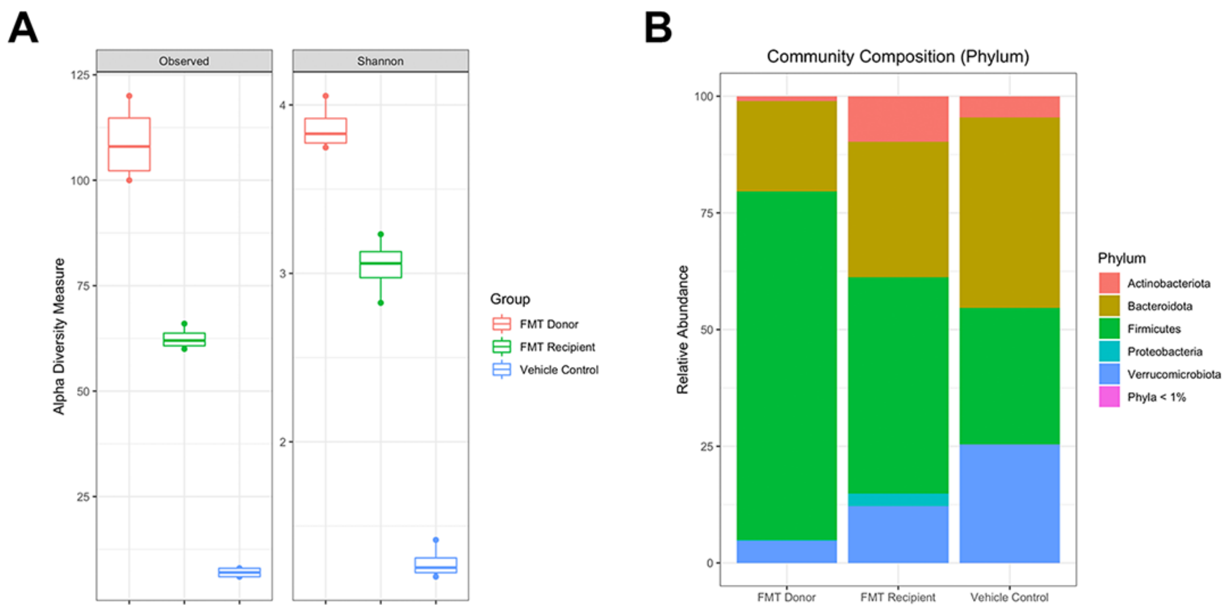


**Supplemental Figure 2:** *Community diversity and composition in the RNA sequencing subset is representative of the overall sample set.* A) Alpha diversity of cecal contents from FMT donor (red), FMT recipient (green), and vehicle control (blue) samples with subsequent RNA sequencing data. FMT recipient diversity was intermediate between FMT donor and vehicle control samples. B) Phylum-level community composition from FMT donor, FMT recipient, and vehicle control samples with subsequent RNA sequencing data.
